# Data mining and experimental approaches to identify combinations of natural herbs against bacterial infections

**DOI:** 10.1101/2022.11.29.518436

**Authors:** Ekansh Mittal, Susan Duncan, Steven Chamberlin

**Affiliations:** Westview High school, Portland, OR, USA; Meadow Park Middle School, Portland, OR, USA; Oregon Health & Science University, Portland, OR, USA

**Author notes:** Correspondence: Ekansh Mittal.

## Abstract

Various studies have identified that natural herbs can be repurposed to treat infectious and bacterial diseases. The purpose of this study is first to test the medicinal value of five herbs including asafoetida, cumin, fenugreek, neem, and turmeric as single agent and in pairs using the bacterial zone of inhibition assay. Second, we used target and network analyses to predict the best combinations. We found that all the herbs as single agent were effective against bacterial infection in the following descending order of efficacy: cumin > turmeric > neem > fenugreek > asafoetida as compared to vehicle (ethanol) treated control. Among all the tested combinations the turmeric and fenugreek combination had the best efficacy in inhibiting the bacterial growth. Next to understand the mechanism of action and to predict the effective combinations among available herbs, we used a data mining and computational analysis approach. Using NPASS, BindingDB, and pathway analysis tools, we identified the bioactive compounds for each herb, then identified the targets for each bioactive compound, and then identified associated pathways for these targets. Then we measured the target/pathway overlap for each herb and identified that the most effective combinations were those which have non-overlapping targets/pathways. For example, we showed as a proof-of-concept that turmeric and fenugreek have the least overlapping targets/pathways and thus is most effective in inhibiting bacteria growth. Our approach is applicable to treat bacterial infections and other human diseases such as cancer. Overall, the computational prediction along with experimental validation can help identify novel combinations that have significant antibacterial activity and may help prevent drug-resistant microbial diseases in human and plants.

## Introduction

The growing concerns related to bacterial and infectious diseases have recently led to the discovery of natural antimicrobials to control microbial diseases. Bacterial diseases include cholera, diphtheria, bacterial meningitis, tetanus, Lyme disease, gonorrhea, and syphilis and tuberculosis. Further, foodborne illnesses are caused by variety of bacteria including *Escherichia coli* (*E. coli*), *Salmonella, Shigella, Norovirus*, and *B. subtilis (Bacillus)[1]*. In fact, recently, *E. coli* O417:H7 caused 58 sicknesses, 22 hospitalizations, 2 cases of advanced liver failure, and 2 deaths in the U.S. and Canada alone [2]. Similarly, tuberculosis, one of the deadliest bacterial infections in the <u>world and a total of 1.6 million people died from TB in 2021[3],</u> without a cure. Clearly these illnesses can cause havoc and there are not many effective cures. Therefore, there is an immediate need to find alternate treatment options. Previous studies show that malaria which is a deadly disease can be cured using plant artemisinin. This shows that plants can cure bacterial and parasitic infections.

Nature is a rich resource of various medicinal plants and food spices. Many ethnic groups for thousands of years have been using these herbs as medicine to treat common health complications. This usage had stimulated the repurposing of assorted herbs to treat various diseases and some herbs have been proven to have antibacterial and anticancer properties [4-7]. Additionally, the natural plant product-based antimicrobials drugs are likely to be effective against multi-drug resistant microbes [8].

In this study we focused to quantify antibacterial activities of asafoetida, cumin, fenugreek, and turmeric. Previous studies have shown that turmeric is very helpful against arthritis. The main bioactive compound is curcumin [9]. Its chemical formula is C_21_H_20_O_6_. Turmeric, in ethanolic extracts has shown activity against *S. epidermidis, Staphylococcus aureus, Klebsiella pneumoniae* and many other diseases [10-15]. Cumin is another herb that is commonly used to improve digestive issues. Its major bioactive compound is an oil substance, cuminaldehyde with the chemical formula of C_10_H_12_O. Cuminaldehyde is 30%-50% of the composition of a cumin seed [16-18]. Cumin had shown activity against food contracted illness such as *Campylobacter jejuni* and *Campylobacter coli* [19]. Asafoetida is mostly used for digestive issues, but has positive effects on the skin and lungs. It has many bioactive compounds. But the one that has the most efficacy against bacterial infection is *Umbelliferone*. The chemical formula of it is C_9_H_6_O_3_ [20]. The extracts of asafoetida have shown activity against acne causing bacteria such as *Propionibacterium acnes* and *Staphylococcus epidermidis*. Fenugreek is well known for its help with digestive issues such as nausea and loss of appetite. Its main bioactive compound is diosgenin and its chemical formula is C_27_H_42_O_3_. The extracts of fenugreek have shown activity against *Staphylococcus* [21]. Neem (Azadirachta indica) is known for its help against skin and eye problems. One if its main compounds is azadirachtin whose chemical formula is C_35_H_44_O_16_ [22] and has medicinal value against *P. gingavalis* (Figure 1).

**Figure 1:**
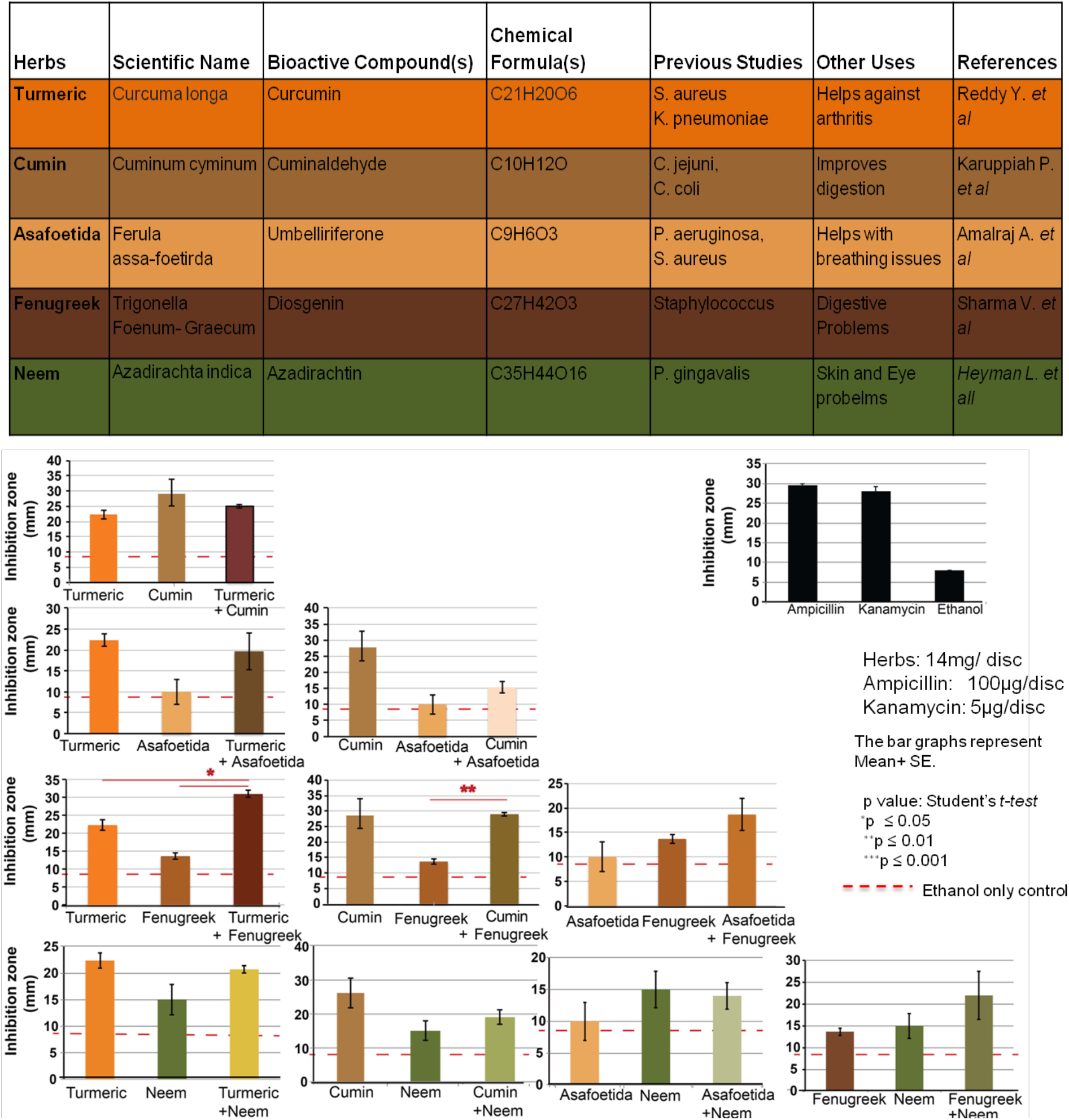
The effect of various herbs prepared in ethanol for testing the anti-bacterial activity against *E. coli* K-12 for establishing zone of inhibition assay. Table 1: The available information and main bioactive compounds for the tested herbs. The experiments were performed in 3 replicates and data is represented as mean± standard deviation (SD) (Related to Supplemental Figure 2 and Supplemental Table 1)

The different herbs have different bioactive compounds that can target different proteins in bacteria to synergize and complement each other’s efficacy. Thus, combining two different herbs may synergize by targeting two different proteins that are important for the survival of bacteria. Therefore, in this research we focused on testing five herbs in combination of two creating 10 different combinations against bacterial growth. However, the experimental testing of all feasible combinations for the available herbs could be time consuming and daunting task. Therefore, in this study we thought to exploit data mining and computational approaches to predict combinations of herbs that are effective for limiting bacterial growth using existing databases. We found that turmeric and fenugreek combination had the best efficacy in inhibiting the bacterial growth and predicted to best combinations using computational approaches. Our approach also has broader implications for diseases such as cancer.

## Materials and Methods

The experiments were performed at the OHSU Primate Research Lab designed specifically for outreach and education programs. We purchased herbs and spices, asafoetida, cumin, fenugreek, and turmeric, and neem from commercial sources tested them against *Escherichia coli (E. coli*) K12. The E.Coli, K12 was purchased by the school and sent directly to the lab.

### Culturing the Escherichia coli K12

*E. coli* K12 was obtained from ATCC as an agar stab. The bacteria cells were cultured overnight at 37°C in Luria-Bertani (LB) broth in a shaking incubator. Different dilution of bacterial culture was plated on nutrient agar plates to determine what dilution of bacteria will give the appropriate bacterial lawn in 22 hours.

### Preparation of the extracts from herbs

The extracts from herbs were prepared by mixing 1 milligram of each herb into 1.5 milliliter of boiling water and 1.5 milliliter of ethanol mix (3 milliliter of liquid). This mixture was vortexed for three days in a cold room which was set to a temperature of 4°C. The extracts were centrifuged at 3,000 rpm for 10 mins and filtered through strainer. All herbs were mixed in 1:1 ratio for creating 6 combinations (Supplemental Figure 1).

### Preparation of antibacterial discs

7 mm discs were prepared from Whatman™ filter paper no 1 using a hole puncher. The discs were autoclaved prior to use and only handled using the autoclaved forceps. Each disc was socked in 42 ml of undiluted extract leading to the concentration of 14 mg/disc.

### Determining the zone of inhibition

*E. coli* K12 bacteria was spreaded across the LB agar plates. Then herb extract socked discs were placed on the agar plate with *E. coli* K12 in 4 replicates. The plates were incubated at 37° C. Several experimental controls were used including 1) extract only control (with no bacteria) 2) solvent only control 3) standard of care antibiotics (Ampicillin: 100ug/disc, as positive controls). After twenty-two hours of incubation the zone of clearance was measured using a ruler in mm.

### Data mining approach for the prediction of effective combinations and impact of herbs on targets and pathways

NPASS (https://bidd.group/NPASS/) was used to identify the bioactive compounds for each herb [23]. Then each compound was matched with a public web-accessible BindingDB database to find potential targets across 30 species [24]. This database allows us to access candidate drug-targets for drug-like molecules and supports medicinal chemistry and drug discovery via literature mining. The target identified using BindingDB data was then matched with Reactome Knowledgebase analysis tool to find potentially interacting pathways [25]. Reactome is an open-source, open access, manually curated and peer-reviewed pathway database. The target and pathway overlaps were identified between two herbs as represented in Venn diagrams. Finally, Stringdb pathway analysis tool (https://string-db.org) was used to identify the effect of herbs on protein-protein interactions and biological processes [26]. STRING is a database of known and predicted protein-protein interactions. The interactions include direct (physical) and indirect (functional) associations; they stem from computational prediction, from knowledge transfer between organisms, and from interactions aggregated from other (primary) databases. Interactions in STRING are derived from five main sources: **(A)** Genomic Context Predictions, **(B)** High-throughput Lab Experiments, **(C)** (Conserved) Co-Expression **(D)** Automated Text mining, and **(E)** Previous Knowledge in Databases. The STRING database currently covers 67.6 million proteins from 14094 organisms. The Stringdb also calculated the false discovery rate (FDR) for finding any association [23].

### Statistical Analysis

Averages and standard errors were calculated by taking into account all 4 replicates. For calculating *P* values, paired, 2-tailed student *t-*test was used for comparing the herbs against the respective solvent control and standard-of-care antibiotic control. Additionally, we calculated the overall significance. To do this we took the *p* value of the herb compared against the solvent control, and divided it by the p value of the herb compared against the antibiotic control. The extracts which have lowest ratio for the p value for solvent to antibiotics would be considered to have highest overall significance (OS) because we wanted herbal extracts to be highly effective compared to no treatments but comparable to the efficacy of antibiotics against bacteria. Then we considered the most effective drugs that has OS value less than 1, the range of 1 to 5 was designated in medium range and OS value above 5 was considered ineffective.

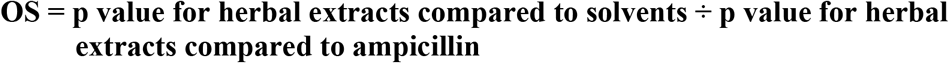

## Results

### Effect of herbs as a signal agent on bacterial growth

We tested asafoetida, cumin, fenugreek, neem, and turmeric herbs against *E. coli* K-12 by preparing ethanolic extracts of each herb. We measured the zone of inhibition which is the area with lesser or no bacteria. The maximum zone of inhibition (ZOI) of a single agent was observed with cumin at 29.0 mm (Figure 1, Supplemental Table 1). The second most was observed with turmeric at 22.33 mm. All singular herbs were effective against bacterial infection (Cumin > Turmeric > Neem > Fenugreek > Asafoetida) compared to vehicle (ethanol) treated control.

### Effect of combination herbs against bacterial growth

We tested each herb in combination and found that multiple combinations are more effective than the single agents. The combination of turmeric and fenugreek had the largest zone of inhibition (ZOI) 31.0+1.0 mm, significantly larger than turmeric and fenugreek singularly which had 22.33±1.45 mm and 13.67<u>+0.88</u> mm ZOI, respectively (Figure 1). Also displayed the second-best overall significance (OS) (Figure 2). The combination of cumin and fenugreek displayed significant zone of inhibition compared to single agent, the highest OS and second-best ZOI with 29+0.58 mm. Interestingly that turmeric and fenugreek, the two of the three best single agents combined had the best combination. But the ZOI was 9.0 mm more than turmeric which was the greater of the two, suggesting a synergistic effect. Fenugreek is also a good herb when combined with cumin. On the other hand, asafoetida was not effective as a single herb but when combined with other herbs it shows reduced or no additional efficacy suggesting antagonistic effects. Of the combinations extracts the order of effectiveness is as follows: Turmeric+Fenugreek > Cumin+Fenugreek > Turmeric + Cumin > Fenugreek+Neem > Turmeric+Neem > Turmeric+Asafoetida > Cumin + Neem > Asafoetida + Fenugreek > Cumin + Asafoetida > Asafoetida + Neem (Figures 1 and 2).

**Figure 2:**
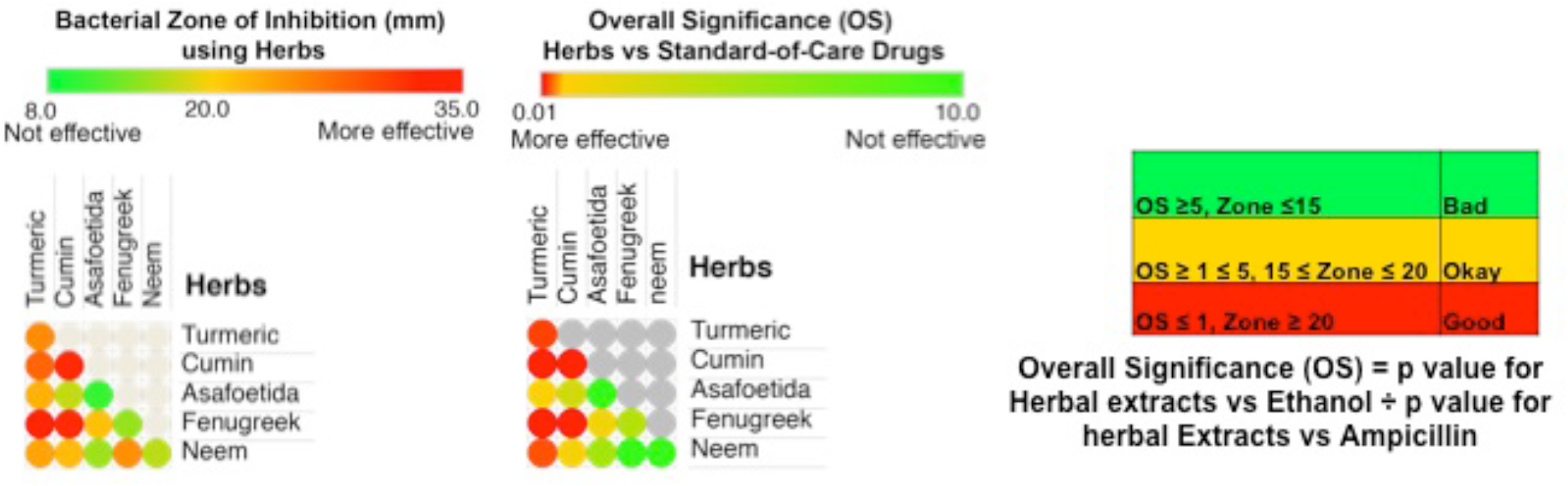
The heatmap representation of bacterial zone of inhibition (ZOI) and overall significance as determined by dividing the p values for herbal extracts compared to solvents with p values for herbal extracts compared to ampicillin. A heat map representing the ZOI is displayed. If the ZOI was >20 mm it was considered an effective inhibition, ZOI range 15 to 20 mm was considered moderate effect, and ZOI <15 mm was considered ineffective against bacterial growth. The most effective herbs were considered those that has overall survival (OS) value <1, the range of 1 to 5 was designated in medium range and OS value >5 was considered ineffective.

### Data curation and computational analysis to find the effective combinations

There are thousands of known herbs consist of various bioactive compounds [27-30]. But if we were to test for example 5,000 herbs and only make combinations of 2, then we have 12,497,500 combinations. Testing of all these combinations won’t be feasible without finding a strategy to prioritize the combinations. Therefore, we established a data mining approach to identify the effective combinations. For this NPASS (http://bidd2.nus.edu.sg/NPASS/about.php)[23] was used to identify the bioactive compounds for each herb (Figure 3). Then each compound was matched with BindingDB to find potential targets across 30 species[24]. The BindingDB data was then matched with Reactome Knowledgebase analysis tool to find potentially interacting pathways[25]. The target and pathway overlaps were identified between two herbs and represented in Venn diagrams. We found that in both turmeric + asafoetida and asafoetida + fenugreek combinations asafoetida had more contribution to overlapping pathways, 72% (79/101) and 63% (70/101) compared to their combination partners with turmeric 12.6% (79/578) and fenugreek 20% (70/345) pathways, respectively in *H. sapiens*. Similarly, the target overlaps for asafoetida with its combination partners turmeric or fenugreek is 62.5% (10 out of 16) (Figure 3). However, the ZOI is better for turmeric (22.33<u>+1.45 mm</u>) or fenugreek (13.67<u>+ 0.88 mm</u>) compared to asafoetida (10.0<u>+3.0 mm</u>) suggesting most of the unique pathways were found for turmeric or fenugreek only in these combinations (Figures 1 and 3). Whereas in turmeric + fenugreek combination, there is the minimal target overlap and broader class of target inhibition. For example, for turmeric only 20% target (29 out of 150) and 46% pathway (257 out of 548) overlaps with fenugreek, offering better effect on bacterial growth inhibition. These results are in consistent with pathway analysis suggest that the turmeric + fenugreek combination inhibits pathways related to SRC kinase and immune function (Figure 3). As a next step to analyze the target protein-protein interactions and network we used Stringdb online tool. This tool allowed us to look at pathway interactions specific to *E. coli* bacteria and human cells. The data suggest that in *E. coli* these herbs inhibit biological processes such as biocarbonate transport and ion transport, probably necessary for survival of the bacterial cells. However, in human cells, these herbs inhibit many diseases related pathways. For example, turmeric and fenugreek inhibits many cancers related pathways such as NF-kb [31], SRC [32], RUNX1[33], and PI3K pathways[34] which are involved in variety of cancer and might also be mutated or have aberrant expression in these cancers (Figure 4). These data suggest that data mining and computational prediction followed by experimental validation may identify the effective combinations in a rapid manner (Figure 4, Supplemental Table 2). This testing can also predict target to use herbs for human, animal, and plant related diseases.

**Figure 3:**
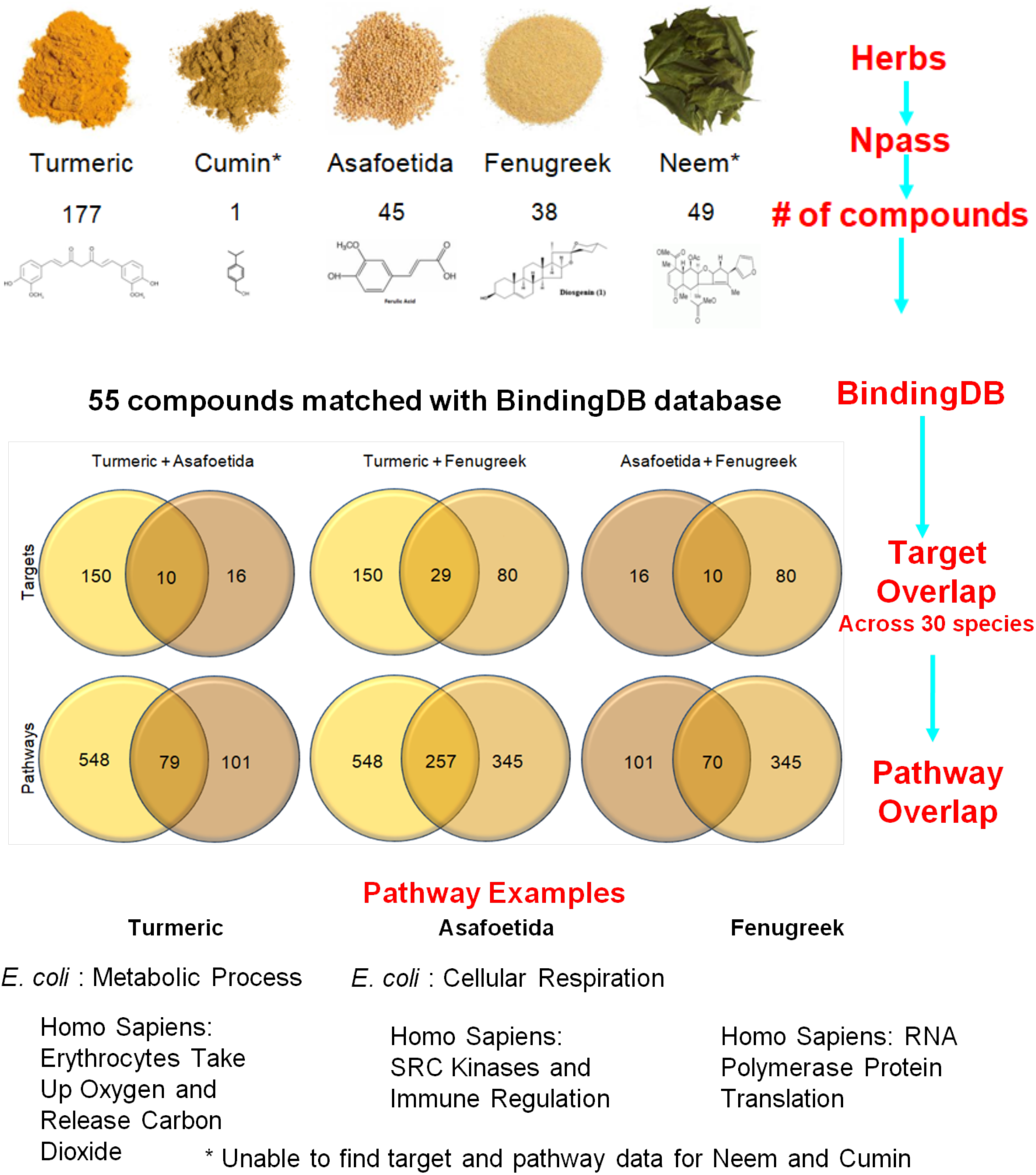
Representation of computationally predicting synergies for the combination of herbs. NPASS was used to identify the bioactive compounds for each herb. Then each compound was matched with binding DB to find potential targets across 30 species. The Binding DB data was then matched with reactome pathway to find potential pathways targets by these herbs. Although we tested herbs in bacteria (*E. coli*), we did pathway analysis for both *E. coli* and human (*Homo sapiens*) to test the broader applicability of the computational approach. Lower panels are Venn diagrams to show an overlap of targets and pathways.

**Figure 4:**
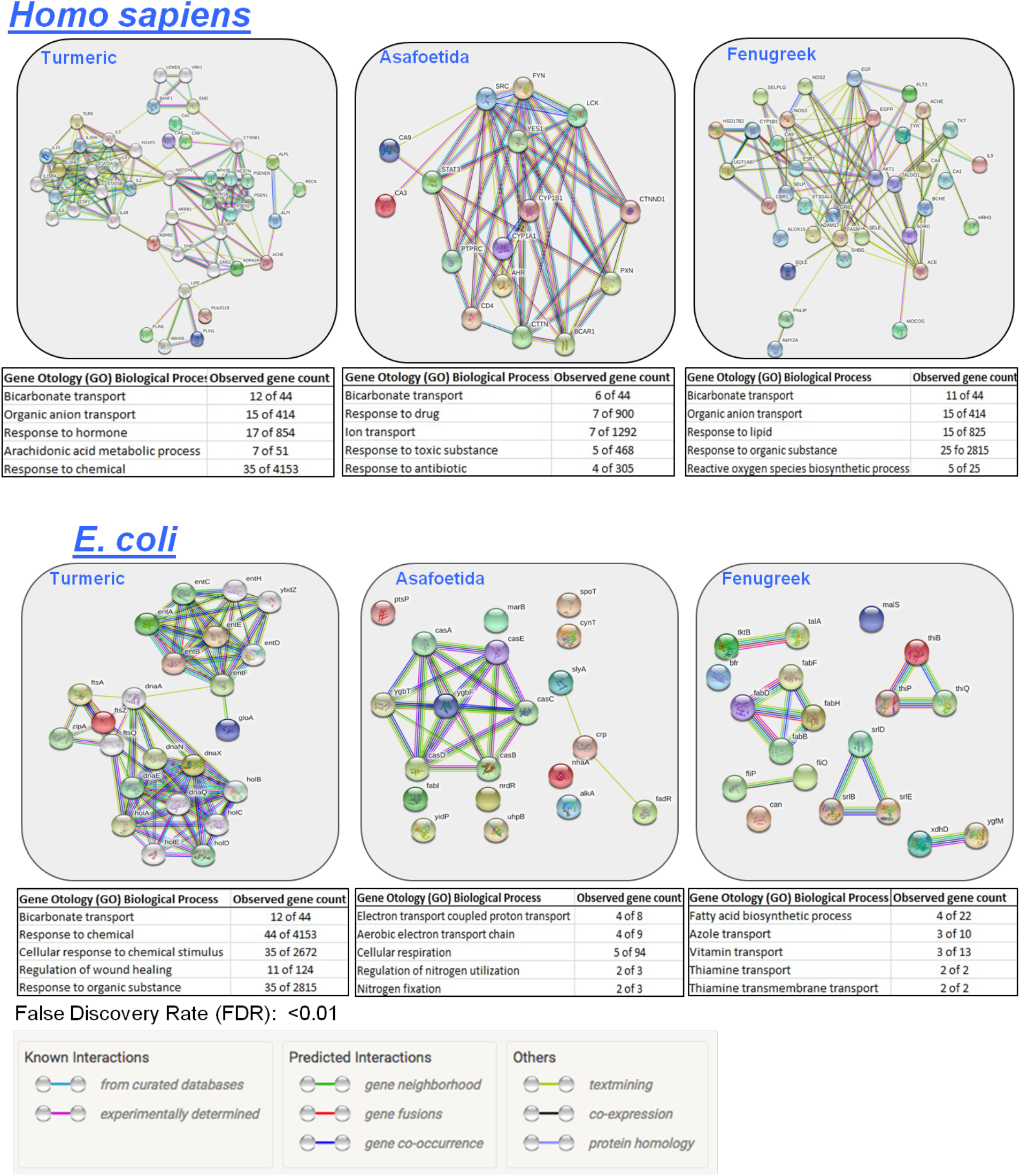
Herb’s target network analysis as identified using string db for the specific organism (*Homo sapiens* or *E. coli*) to gain additional knowledge of target protein interactions and pathway network that might be impacted by individual herbs. The example of pathways and protein-protein interactions impacted by each herb is shown for both human and *E. coli*. Cumin and Neem has no target data available. The table underneath of each pathway shows that which biological processes will be impacted by these herbs. These biological processes are defined based on Gene Otology term (GO). False discovery rate (FDR) for each of the biological processes is less than 0.01.

## Discussion

We found that herbal extracts in different combinations are more effective in suppressing bacterial growth compared to singular herbs. As a single agent, cumin and turmeric remained the best (cumin > turmeric > neem > fenugreek > asafoetida) in inhibiting bacterial growth with zone of inhibition of 29.0 + 4.51 and 22.33+ 1.45, respectively. Turmeric and fenugreek combination had a zone of inhibition of 31.0+1.0 mm which is 2.3 fold bigger than fenugreek which had a zone of inhibition of 13.67+0.88 mm and is 1.4 times more potent than turmeric which had a zone of inhibition of 22.33+1.45 mm. Of the combinations extracts the order of effectiveness is as follows: turmeric+fenugreek > cumin + fenugreek > turmeric + cumin > fenugreek + neem > turmeric + neem > turmeric + asafoetida > cumin + neem > asafoetida + fenugreek > cumin + asafoetida > asafoetida + neem. Though turmeric and fenugreek and cumin and fenugreek both combinations showed similar zone of inhibition, the turmeric and fenugreek were the only combination that showed significantly more effective compared to the single treatments in inhibiting the growth of *E. coli* K 12. Most of the other combinations seemed to show average inhibition of the two combined herbs or similar to the effect of single herb, suggesting only one herb is effective. Our results suggest that certain combinations of herbs may work better than single agents in eliminating bacterial infections.

However, there are thousands known herbs and there could be billion different combinations feasible. If we were to test all of them one by one, it would take, decades, maybe even centuries. Therefore, we need a systematic way to predict these combinations before experimental validations. Our data mining approach suggests that computational or machine learning approaches can help achieving these at faster rate. Then allow us to test these selected combinations in wet lab settings. Computational analysis was also helpful to translate these findings to human disease as reactome pathway and Stringdb protein-protein interactions and gene ontology analyses identified that some of the combinations that are effective against bacterial growth could also be effective in targeting the pathway that are associated with cancer and other diseases.

In the future, it will be important to optimize the methods for the extraction of the bioactive compounds from these herbs as it is possible that all the bioactive compounds are not fully extracted by our method. Further, it will be critical to test the medicinal values of purified bioactive compounds for potency as whole plant extract might have multiple bioactive compounds and many interactions might be occurring within bioactive compounds whole plant extract[35-37]. Additionally, whole plant extracts are often metabolized to other compounds in an organism, so you maybe be getting a very different compound exposure for a microbe[38]. Further, the current databases have limited information for many of these herbs and requiring multiple database mining. Thus, combining all these information in one database will improve the efficiency of this curation. We also do not fully understand the mechanisms by which these combinations are effective in blocking bacterial growths. This information may help finding the synergistic herbs. These combinations of herbs may have broader implications against infectious disease and cancers in humans, plants, and animals [39].

## Author Contributions

EM planned and performed the experiment, did data analysis, wrote the research paper. SD and SC provided resources and mentorship. All the authors edited the manuscript.

## Acknowledgments

We thank Dr. Diana Gordon (Oregon Health and Science University Primate Center) and Aparna Govindan (Oregon Health and Science University Primate Center) to provide all the resources for performing experiments and mentorship. We also thank Beaverton school district science fair, Broadcom MASTERS, and 3M young scientist challenge to support and encouragement.

## References

1. Ashwathi P. 14 Major Diseases Caused by Bacteria | Microbiology.

2. McKeen A. E. coli outbreak leaves two dead — one in Canada, one in the U.S. TORONTO STAR, 2018.

3. WHO. Tuberculosis. World Health Organization, 2022.

4. El-Ghorab AH, Nauman M, Anjum FM, Hussain S, Nadeem M. A comparative study on chemical composition and antioxidant activity of ginger (Zingiber officinale) and cumin (Cuminum cyminum). J Agric Food Chem 2010; 58:8231–7.

5. Ivanovic M, Makoter K, Islamcevic Razborsek M. Comparative Study of Chemical Composition and Antioxidant Activity of Essential Oils and Crude Extracts of Four Characteristic Zingiberaceae Herbs. Plants (Basel) 2021; 10.

6. M AA-T, Rastall B, I MA-R. Chemical Composition, Cytotoxic, Apoptotic and Antioxidant Activities of Main Commercial Essential Oils in Palestine: A Comparative Study. Medicines (Basel) 2016; 3.

7. Sepahpour S, Selamat J, Abdul Manap MY, Khatib A, Abdull Razis AF. Comparative Analysis of Chemical Composition, Antioxidant Activity and Quantitative Characterization of Some Phenolic Compounds in Selected Herbs and Spices in Different Solvent Extraction Systems. Molecules 2018; 23.

8. Karuppiah P, Rajaram S. Antibacterial effect of Allium sativum cloves and Zingiber officinale rhizomes against multiple-drug resistant clinical pathogens. Asian Pac J Trop Biomed 2012; 2:597–601.

9. Chaitanya BV, Somisetty KV, Diwan A, Pasha S, Shetty N, Reddy Y, et al. Comparison of Antibacterial Efficacy of Turmeric Extract, Morinda Citrifolia and 3% Sodium Hypochlorite on Enterococcus faecalis: An In-vitro Study. J Clin Diagn Res 2016; 10:ZC55–ZC57.

10. Elgamily H, Safy R, Makharita R. Influence of Medicinal Plant Extracts on the Growth of Oral Pathogens Streptococcus Mutans and Lactobacillus Acidophilus: An In-Vitro Study. Open Access Maced J Med Sci 2019; 7:2328–2334.

11. Guru SR, Reddy KA, Rao RJ, Padmanabhan S, Guru R, Srinivasa TS. Comparative evaluation of 2% turmeric extract with nanocarrier and 1% chlorhexidine gel as an adjunct to scaling and root planing in patients with chronic periodontitis: A pilot randomized controlled clinical trial. J Indian Soc Periodontol 2020; 24:244–252.

12. Patil V, Akal N, Biradar S, Ratnakar P, Rairam S, Batta O. Comparative evaluation of antimicrobial efficacy of mushroom, Aloe vera, and Curcuma longa with calcium hydroxide as an intracanal medicament against Enterococcus faecalis: An in vitro study. J Conserv Dent 2022; 25:415–419.

13. Prabhakar A, Taur S, Hadakar S, Sugandhan S. Comparison of Antibacterial Efficacy of Calcium Hydroxide Paste, 2% Chlorhexidine Gel and Turmeric Extract as an Intracanal Medicament and their Effect on Microhardness of Root Dentin: An in vitro Study. Int J Clin Pediatr Dent 2013; 6:171–7.

14. Singh V, Gupta A, Verma UP, Mishra T, Pal M. An evaluation of the efficacy of ethanolic extract of Nigella sativa L. (Kalonji) on the clinical parameters of moderatetosevere gingivitis: A splitmouth clinical study. Ayu 2019; 40:152–158.

15. Stoyell KA, Mappus JL, Gandhi MA. Clinical efficacy of turmeric use in gingivitis: A comprehensive review. Complement Ther Clin Pract 2016; 25:13–17.

16. Bettaieb I, Bourgou S, Sriti J, Msaada K, Limam F, Marzouk B. Essential oils and fatty acids composition of Tunisian and Indian cumin (Cuminum cyminum L.) seeds: a comparative study. J Sci Food Agric 2011; 91:2100–7.

17. Alimohamadi K, Taherpour K, Ghasemi HA, Fatahnia F. Comparative effects of using black seed (Nigella sativa), cumin seed (Cuminum cyminum), probiotic or prebiotic on growth performance, blood haematology and serum biochemistry of broiler chicks. J Anim Physiol Anim Nutr (Berl) 2014; 98:538–46.

18. Bettaieb I, Bourgou S, Wannes WA, Hamrouni I, Limam F, Marzouk B. Essential oils, phenolics, and antioxidant activities of different parts of cumin (Cuminum cyminum L.). J Agric Food Chem 2010; 58:10410–8.

19. Derakhshan S, Sattari M, Bigdeli M. Effect of cumin (Cuminum cyminum) seed essential oil on biofilm formation and plasmid Integrity of Klebsiella pneumoniae. Pharmacogn Mag 2010; 6:57–61.

20. Amalraj A, Gopi S. Biological activities and medicinal properties of Asafoetida: A review. J Tradit Complement Med 2017; 7:347–359.

21. Al-Timimi LAN. Antibacterial and Anticancer Activities of Fenugreek Seed Extract. Asian Pac J Cancer Prev 2019; 20:3771–3776.

22. Alzohairy MA. Therapeutics Role of Azadirachta indica (Neem) and Their Active Constituents in Diseases Prevention and Treatment. Evid Based Complement Alternat Med 2016; 2016:7382506.

23. Zeng X, Zhang P, He W, Qin C, Chen S, Tao L, et al. NPASS: natural product activity and species source database for natural product research, discovery and tool development. Nucleic Acids Res 2018; 46:D1217–D1222.

24. Liu T, Lin Y, Wen X, Jorissen RN, Gilson MK. BindingDB: a web-accessible database of experimentally determined protein-ligand binding affinities. Nucleic Acids Res 2007; 35:D198–201.

25. Gillespie M, Jassal B, Stephan R, Milacic M, Rothfels K, Senff-Ribeiro A, et al. The reactome pathway knowledgebase 2022. Nucleic Acids Res 2022; 50:D687–D692.

26. Szklarczyk D, Gable AL, Lyon D, Junge A, Wyder S, Huerta-Cepas J, et al. STRING v11: protein-protein association networks with increased coverage, supporting functional discovery in genome-wide experimental datasets. Nucleic Acids Res 2019; 47:D607–D613.

27. Kiroglu O, Berktas F, Khan Z, Dagkiran M, Karatas Y. Self-medication practices with conventional and herbal drugs among ear, nose, and throat patients. Rev Assoc Med Bras (1992) 2022; 68:1416–1422.

28. Booncharoen P, Boonchai W, Akarasereenont P, Tripatara P. A comparative study of chemical constituents and safety of Thai herbal medicated oil formula and traditional medicated oil. J Complement Integr Med 2021.

29. Dechayont B, Phuaklee P, Chunthorng-Orn J, Juckmeta T, Prajuabjinda O, Jiraratsatit K. Antibacterial, anti-inflammatory and antioxidant activities of Mahanintangtong and its constituent herbs, a formula used in Thai traditional medicine for treating pharyngitis. BMC Complement Med Ther 2021; 21:105.

30. Wei X, Zhao Z, Zhong R, Tan X. A comprehensive review of herbacetin: From chemistry to pharmacological activities. J Ethnopharmacol 2021; 279:114356.

31. Alaaeldin R, Ali FEM, Bekhit AA, Zhao QL, Fathy M. Inhibition of NF-kB/IL-6/JAK2/STAT3 Pathway and Epithelial-Mesenchymal Transition in Breast Cancer Cells by Azilsartan. Molecules 2022; 27.

32. Shao W, Liu L, Zheng F, Ma Y, Zhang J. The potent role of Src kinase-regulating glucose metabolism in cancer. Biochem Pharmacol 2022; 206:115333.

33. Lin TC. RUNX1 and cancer. Biochim Biophys Acta Rev Cancer 2022; 1877:188715.

34. Madsen RR. Principles of Pi3k Biology and Its Role in Lymphoma. In: O’Conner Oa, Ansell SM, Seymour JF, editors. Targeting Oncogenic Drivers and Signaling Pathways in Hematologic Malignancies: From Concept to Practice. Hoboken (NJ), 2023.

35. Gomathi D, Kalaiselvi M, Ravikumar G, Devaki K, Uma C. GC-MS analysis of bioactive compounds from the whole plant ethanolic extract of Evolvulus alsinoides (L.) L. J Food Sci Technol 2015; 52:1212–7.

36. Kennedy DO, Wightman EL. Herbal extracts and phytochemicals: plant secondary metabolites and the enhancement of human brain function. Adv Nutr 2011; 2:32–50.

37. Vaou N, Stavropoulou E, Voidarou CC, Tsakris Z, Rozos G, Tsigalou C, et al. Interactions between Medical Plant-Derived Bioactive Compounds: Focus on Antimicrobial Combination Effects. Antibiotics (Basel) 2022; 11.

38. Salem MA, Perez de Souza L, Serag A, Fernie AR, Farag MA, Ezzat SM, et al. Metabolomics in the Context of Plant Natural Products Research: From Sample Preparation to Metabolite Analysis. Metabolites 2020; 10.

39. Khan T, Ali M, Khan A, Nisar P, Jan SA, Afridi S, et al. Anticancer Plants: A Review of the Active Phytochemicals, Applications in Animal Models, and Regulatory Aspects. Biomolecules 2019; 10.

