## Supplemental Figures 1-2 and Table 1 for "Data mining and experimental approaches to identify combinations of natural herbs against bacterial infections" (from another version of this preprint: version 1, 2022-11-29)

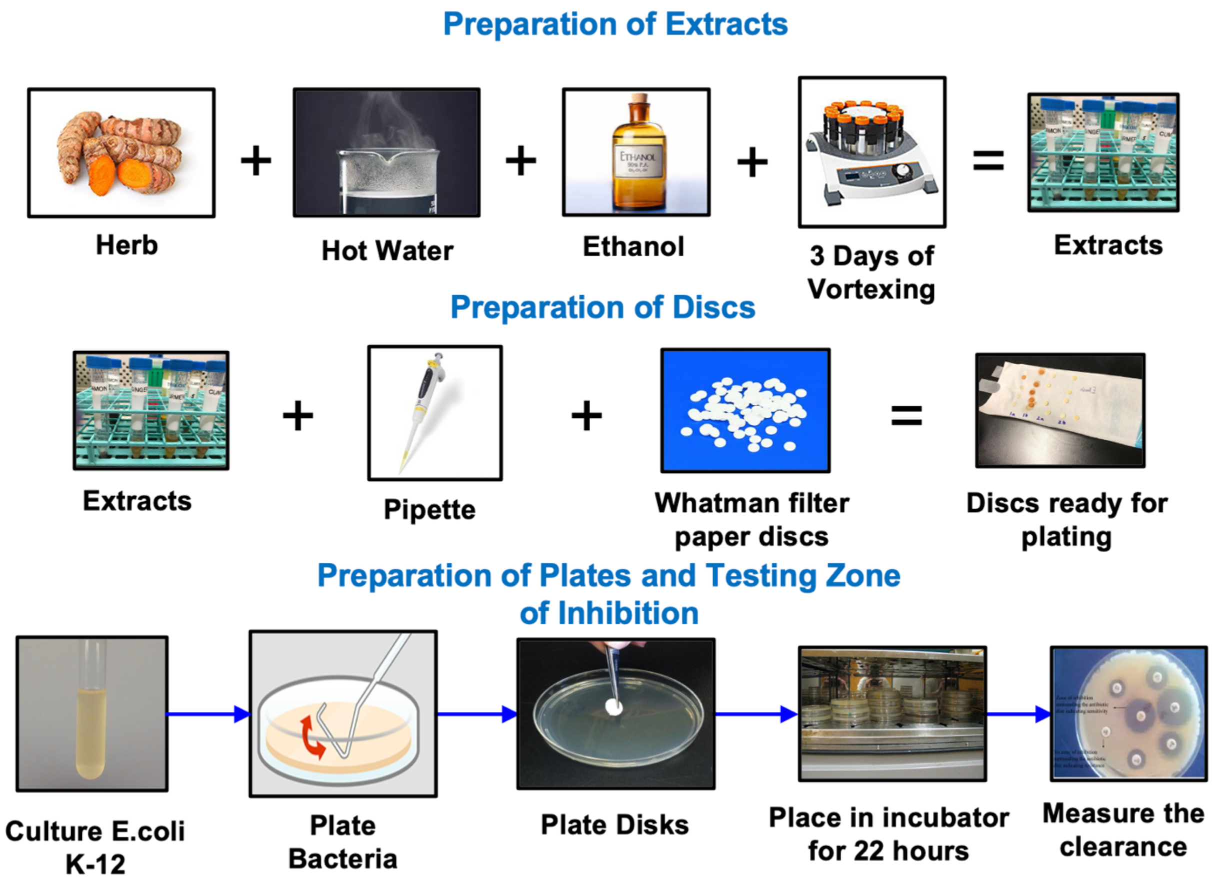


**Supplemental Figure 1: Method for preparing extracts from herbs and testing zone of inhibition using E coli K-12 bacteria.**

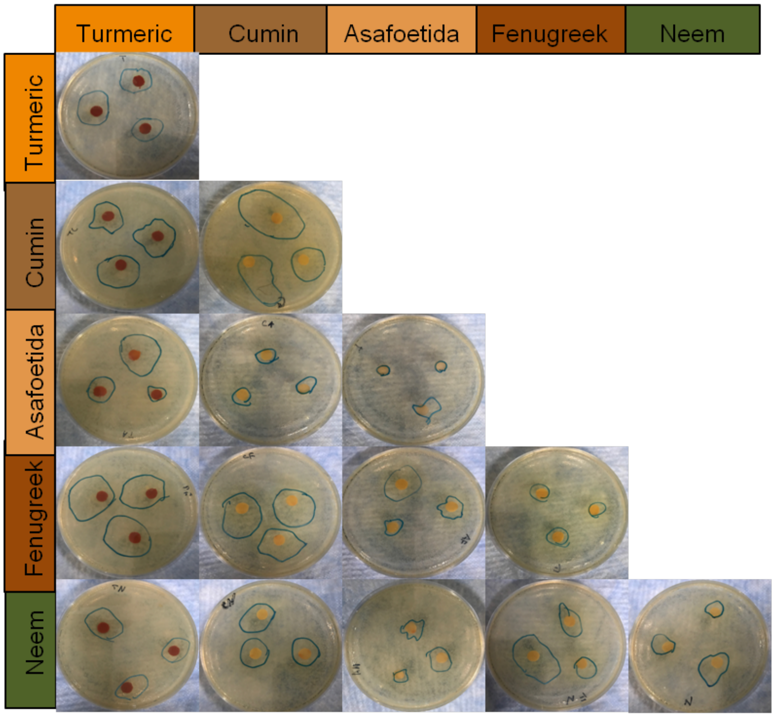


**Supplemental Figure 2: The agar plates showing the effect of various herbs on the anti-bacterial activity of *E. coli* K-12.** Each disc has herbal extract equivalent to 14 mg per disc. Each herb was tested in triplicate in combination as well as single treatment along with the standard-of care treatment drugs.

lue

| | Average | SE | Ethanol Vs Herb | Combo vs Single | Treatment vs Amp | OS |
|---|---|---|---|---|---|---|
| | | | **P Values** | | | |
| Turmeric | 22.33 | 1.45 | 0.010 | 0.184 | 0.053 | 0.19 |
| Cumin | 29.00 | 4.51 | 0.043 | 0.508 | 0.892 | 0.05 |
| Turmeric+Cumin | 25.00 | 0.58 | 0.001 | | 0.020 | 0.06 |
| Turmeric | 22.33 | 1.45 | 0.010 | 0.601 | 0.053 | 0.19 |
| Asafoetida | 10.00 | 3.00 | 0.574 | 0.237 | 0.028 | 20.82 |
| Turmeric+Asafoetida | 19.67 | 4.41 | 0.118 | | 0.149 | 0.79 |
| Turmeric | 22.33 | 1.45 | 0.010 | 0.043 | 0.053 | 0.19 |
| Fenugreek | 13.67 | 0.88 | 0.023 | 0.011 | 0.005 | 4.52 |
| Turmeric+Fenugreek | 31.00 | 1.00 | 0.002 | | 0.270 | 0.01 |
| Turmeric | 22.33 | 1.45 | 0.010 | 0.199 | 0.053 | 0.19 |
| Neem | 15.00 | 2.89 | 0.136 | 0.199 | 0.030 | 4.53 |
| Turmeric+Neem | 20.67 | 0.67 | 0.003 | | 0.012 | 0.23 |

| | Average | SE | Ethanol vs Herb | Combo vs Single | Treatment vs Amp | OS |
|---|---|---|---|---|---|---|
| | | | **P Values** | | | |
| Cumin | 29.00 | 4.51 | 0.043 | 0.098 | 0.892 | 0.05 |
| Asafoetida | 10.00 | ## | 0.574 | 0.119 | 0.028 | 20.82 |
| Cumin+Asafoetida | 15.33 | 1.76 | 0.053 | | 0.019 | 2.74 |
| Cumin | 29.00 | 4.51 | 4.509 | 1.000 | 0.892 | 5.06 |
| Fenugreek | 13.67 | ## | 0.023 | 0.006 | 0.005 | 4.52 |
| Cumin+Fenugreek | 29.00 | ## | 0.001 | | 0.184 | 0.00 |
| Cumin | 29.00 | 4.51 | 0.043 | 0.063 | 0.892 | 0.05 |
| Neem | 15.00 | ## | 0.136 | 0.195 | 0.030 | 4.53 |
| Cumin+Neem | 19.00 | ## | 0.034 | | 0.034 | 0.99 |

| | Average | SE | Ethanol vs Herb | Combo vs Single | Treatment vs Amp | OS |
|---|---|---|---|---|---|---|
| | | | **P Values** | | | |
| Asafoetida | 10.00 | 3.00 | 0.574 | 0.274 | 0.028 | 20.82 |
| Fenugreek | 13.67 | 0.88 | 0.023 | 0.286 | 0.005 | 4.52 |
| Asafotida+Fenugeel | 18.67 | 3.28 | 0.083 | | 0.069 | 1.20 |
| Asafoetida | 10.00 | 3.00 | 0.574 | 0.472 | 0.028 | 20.82 |
| Neem | 15.00 | 2.89 | 0.136 | 0.858 | 0.030 | 4.53 |
| Asafoetida+Neem | 14.00 | 2.08 | 0.102 | | 0.021 | 4.76 |

| | Average | SE | Ethanol vs Herb | Combo vs Single | Treatment vs Amp | OS |
|---|---|---|---|---|---|---|
| | | | **P Values** | | | |
| Fenugreek | 13.67 | 0.88 | 0.023 | 0.220 | 0.005 | 4.01 |
| Neem | 15.00 | 2.89 | 0.136 | 0.210 | 0.030 | 13.75 |
| Fenugreek+Neem | 22.00 | 5.51 | 0.126 | 0.297 | | 18.53 |

**Supplemental Table 1: The table showing zone of inhibition (ZOI), p values and Overall significance (OS) values for each herb and its combinations.**
